# Phytoremediation of EMS Induced *Brassica juncea* Heavy Metal Hyperaccumulator Genotypes

**DOI:** 10.1101/2021.12.26.474158

**Authors:** Shema Halder, Apurba Anirban

## Abstract

Buriganga, an economically important river of Dhaka, Bangladesh, is highly polluted by different toxic heavy metals. In this study, phytoremediation of EMS induced Indian mustard (*Brassica juncea* L) genotypes against three pollutants *viz*. lead (Pb), chromium (Cr) and cadmium (Cd) of Buriganga riverbank soil was assessed in field condition. Among 1-, 2- and 3% EMS induced genotypes, better seed germination rate, germination speed and plant survival rate were observed in 1% EMS induced genotype, BE21. The highest concentration of Pb, Cr and Cd were also obtained in the leaf of BE21 genotype and therefore was considered as a super-hyperaccumulator genotype. Concentration of Pb in the next generation of this genotype was approximately two-fold higher in the root (91.53±6.59 mg/kg dry weight, DW); three-fold higher in the shoot (33.31±1.01 mg/kg DW) and leaf (28.35±3.61 mg/kg DW), and more in the fruit (5.59±0.93 mg/kg DW) than the control. Concentration of Cr was approximately two-fold in the root (57.02±3.24 mg/kg DW), shoot (18.51±1.36 mg/kg DW) and leaf (14.98±2.01 mg/kg DW), and more in the fruit (6.15±1.92 mg/kg DW) of BE21 genotype compared to the control. Cd concentration was more in the root (1.96±0.92 mg/kg DW), leaf (0.52±0.32 mg/kg DW) and fruit (0.19±0.02 mg/kg DW) and less in the shoot (0.19±0.01 mg/kg DW) of BE21 genotype than the control. Root, shoot, leaf and fruit of BE21 altogether accumulated 98-, 73- and 87% Pb, Cr and Cd, respectively and can thus be utilized to remove heavy metals of Buriganga River. As like root, shoot and leaf, fruit also accumulated heavy metals; hence those plants which are used in phytoremediation should not be used as food or fodder. To the best of our knowledge, this is the first report of developing EMS induced hyperaccumulator genotype of *B. juncea* for phytoremediation of Buriganga riverbank soil of Bangladesh.

## INTRODUCTION

Increased metal concentrations have adverse effects on various living organisms (Ahmad et al., 2011; John et al., 2009). Therefore, various techniques were developed for the remediation of contaminated soils. Phytoremediation is a new but promising technology for cleanup of polluted sites and also requires less investment than physicochemical technologies (Garbisu and Alkorta, 2001; McGrath et al., 2001). In many countries, the polluted soil is treated with physical, chemical and thermal processes which are very much costly. With those processes, polluted soil site cleanup requires 0.6–2.5 million/m3 (McIntyre, 2003); whereas the same area requires USD $37.7 when plants with deep roots are used for phytoremediation (Wan et al., 2016). As such, phytoremediation technology is cost effective, eco-friendly and thus alternative to traditional methods.

For the study of phytoremediation, it is important to know which plants are suitable to be considered as hyperaccumulators (Rodriguez et al., 2005). Several investigations were performed using Indian mustard (Choudhury et al., 2016; Ishikawa et al., 2006; John et al., 2009; Salido et al., 2003), rice (Hussain et al., 2021; Murakami et al., 2009; Payus and Talip, 2014; Satpathy et al., 2014; Zakaria et al., 2021), wheat (Chandra et al., 2009; Liu et al., 2009), maize (Wang et al., 2002) etc. to assess their capacity to remediate pollutants. Although different types of plants have been used as heavy metal hyperaccumulators, however wild/cultivated plants those are used in phytoremediation are not able to accumulate much heavy metals. As such, genotypes of various plant species, for example tomato, Arabidopsis, barley etc. induced with chemical agents were identified with better efficiencies regarding phytoremediation (Raskin et al., 1997). Ethyl methane sulfonate (EMS) is one of the most effective and powerful chemicals that could be used to produce hyperaccumulator genotypes. Improvement of plants by genetic modifications also has beneficial impacts on phytoremediation of metal-polluted soils. Researchers induced mutation by EMS in *Brassica rapa* (Mizuno et al., 2014) and in *Brassica juncea* (Ibrahim IA, 2009) to select highly tolerant plants to some heavy metals.

Different waste materials and pollutants containing heavy metals are disposed into Buriganga river of Dhaka city, the capital of Bangladesh. Different contaminants can easily migrate to the environment and thus create contamination of the ecosystems (Gaur, 2004). In addition, contaminants like heavy metals cause toxic effects (Shukla and Singhal, 1984), oxidative stress (Ahmad et al., 2011) to plants, and have impact on nutrition and metabolism of plants (John et al., 2009). It is difficult to remediate the pollutants of Buriganga River by physical, chemical and thermal processes, which require huge cost. In this regard, EMS induced super-accumulator genotypes may be the best alternative for phytoremediation of polluted site of Buriganga River. In addition, *Brassica juncea* is a suitable plant for phytoremediation (Turan and Esringti, 2007), genetic approach to produce hyperaccumulator genotypes of *B. juncea* using EMS is therefore essential. Considering all of the aforementioned aspects, objectives of this research project are— a) induction of mutation in *B. juncea* using EMS as a chemical mutagen, b) selection of heavy metal tolerant plants, and c) assessment of phytoremediation potential of EMS induced hyperaccumulator and super-hyperaccumulator genotypes, respectively.

## MATERIALS AND METHODS

### Experimental material

Indian mustard (*Brassica juncea* L) seeds (Fig 1A) were collected from local market of Dhaka city and used as a basic genetic material for this study. Assessment of phytoremediation potential was conducted after EMS treatment, followed by heavy metal treatment with lead (Pb), chromium (Cr) and cadmium (Cd). The uppermost soil layer of Buriganga riverside containing waste materials and heavy metals (Fig 1B) was used as a heavy metal source. Heavy metal phytoremediation was performed in two steps. At first, 1-, 2-, and 3% EMS were applied on *B. juncea* seeds, then hyperaccumulator genotypes were selected, and finally phytoremediation of 1-, 2-, and 3% EMS treated 8-week-old leaves of *B. juncea* were performed. In the next step, super-hyperaccumulator genotype was treated with heavy metal polluted soil and phytoremediation of root, shoot, leaf and fruit (seeds) was conducted at 12-week planting period (at full maturity stage).

**Fig 1.**
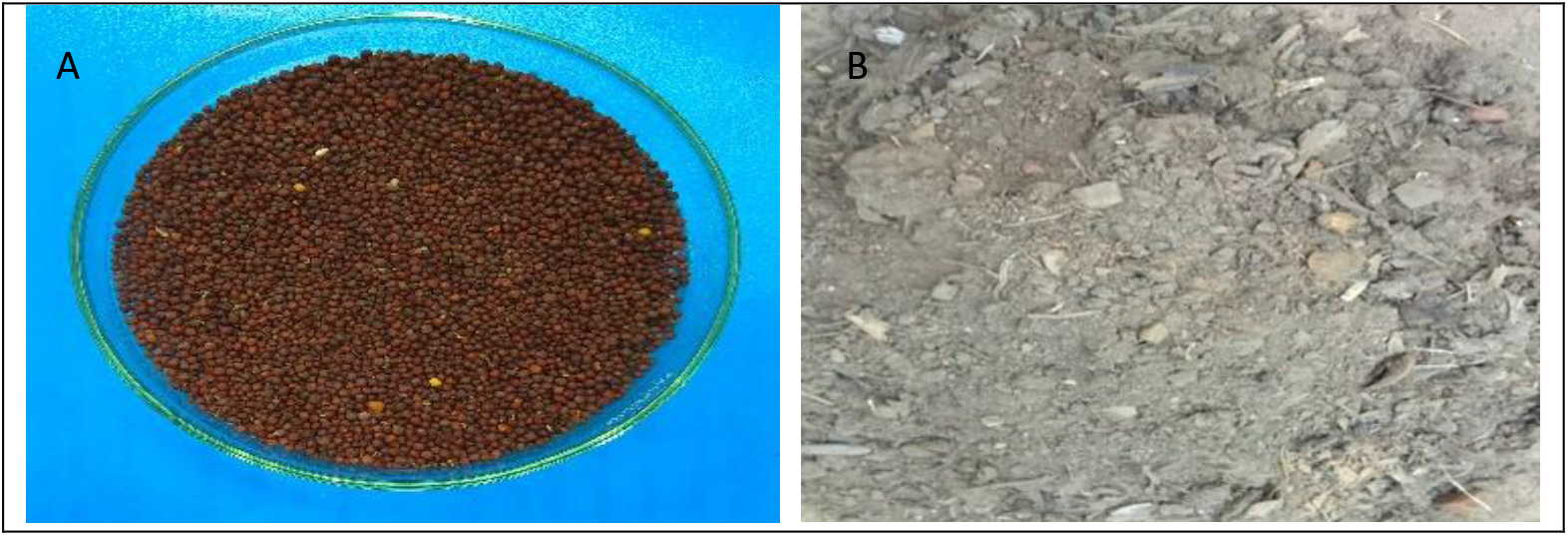
A) Seed samples of *Brassica juncea;* B) Polluted surface soil of Buriganga riverbank.

### EMS Treatment

EMS was used to produce hyperaccumulator genotypes. For the preparation of 1-, 2-, and 3% EMS solution 0.05 ml, 0.1 ml and 0.15 ml EMS, respectively were taken by pipette and added into 5ml *DH_2_O* in three separate beakers. For each percentage of solution 0.25g seeds were taken using an electric balance for EMS treatment. Seeds were soaked in distilled water for 8 h/ overnight at room temperature before EMS treatment. In the following morning, seeds were soaked in different percentage of EMS solution and the mixture was incubated at room temperature for 4 h with gentle stirring. After 4 h, seeds were thoroughly washed in running water for 10 min. EMS treated seeds were grown immediately in the polythene bags filled with polluted soil having heavy metals in the garden of the Department of Botany of Jagannath University, Dhaka. This experiment was conducted between November 2017 and January 2018.

### Heavy metal treatment

Polluted Buriganga riverbank soil was used as a source of heavy metals (Pb, Cr and Cd). Soil containing heavy metals was in heterogeneous condition. Therefore, for the preparation of soil at first soil was mixed properly with the help of a spade. Soil containing waste material *i.e*., plastic element, brickbat, small stone and some other unnecessary things were removed. Small polythene bags were filled with soil and a pore was created at the bottom of each bag for passing extra water. EMS treated seeds were grown immediately in the prepared bags in the garden. For proper growth of plants, weed eradication and irrigation were undertaken properly. No fertilizer was applied for the growth of the plants.

### Selection of heavy metal tolerant plants

For phytoremediation study, hyperaccumulator genotypes from 1-, 2-, and 3% EMS treated *B. juncea* plants were selected based on seed germination rate, germination speed and plant survival rate.

#### a) Seed germination rate

After EMS treatment, seeds were sown in polythene bags having heavy metal polluted soil and their germination status was observed up to one week. Seed germination rate was calculated by the following formulae-

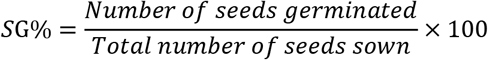

#### b) Germination speed

The germination speed of 1-, 2-, and 3% EMS induced *B. juncea* was calculated by the following formulae as described earlier (Maguire, 1962)—

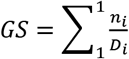

n_i_ =number of seeds germinated on i^th^ day.

D_i_ =number of days from the start of the experiment.

#### c) Plant survival rate

Seedlings those were sown in different polythene bags filled with heavy metal polluted soil were observed up to four weeks of seed sowing and plant survival rate was then calculated by the following formulae—

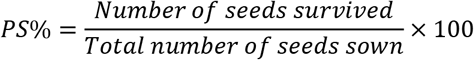

### Assessment of Phytoremediation

Leaf samples from selected plants were collected after eight weeks of 1-, 2-, and 3% EMS induced seed sowing and heavy metal phytoremediation was performed. For super-hyperaccumulator genotype, after 12 weeks of seed sowing root, shoot, leaf and fruit (seeds) samples were collected and heavy metal phytoremediation was conducted.

#### a) Collection of samples

Polluted soil of Buriganga riverbank and plant samples grown in the polluted soil were collected and wrapped with aluminum foil and tagged with small piece of paper. Pb, Cr and Cd concentrations in the leaf samples of 1-, 2-, and 3% EMS induced *B. juncea* were determined by Atomic Absorption Spectrometry. For super-hyperaccumulator genotype, root, shoot, leaf and fruit (seeds) samples were used for assessment of phytoremediation.

#### b) Sample preparation

At first, samples of 1-, 2-, and 3% EMS induced *B. juncea* were washed by running tap water. Samples were further washed by distilled water and then were oven dried. Samples were then cut into small pieces and placed on the Petri dishes (Fig 2A). Approximately 0.5 g samples of each of 1-, 2-, and 3% EMS induced *B. juncea* were measured and placed in the beakers (Fig 2B). For the digestion of plant samples 10 ml conc. *HNO_3_* (Nitric acid) was added with each sample. Then samples were placed on the hot plate. By slow boiling and evaporation process samples became concentrated (Fig 2C). Required deionized water was added with the concentrated samples and poured into 25ml volumetric flask to prepare 25 ml solution. Samples were filtrated with whatman no. 42 filter paper (Fig 2D) and were stored in plastic bottles for metal analysis.

**Fig 2.**
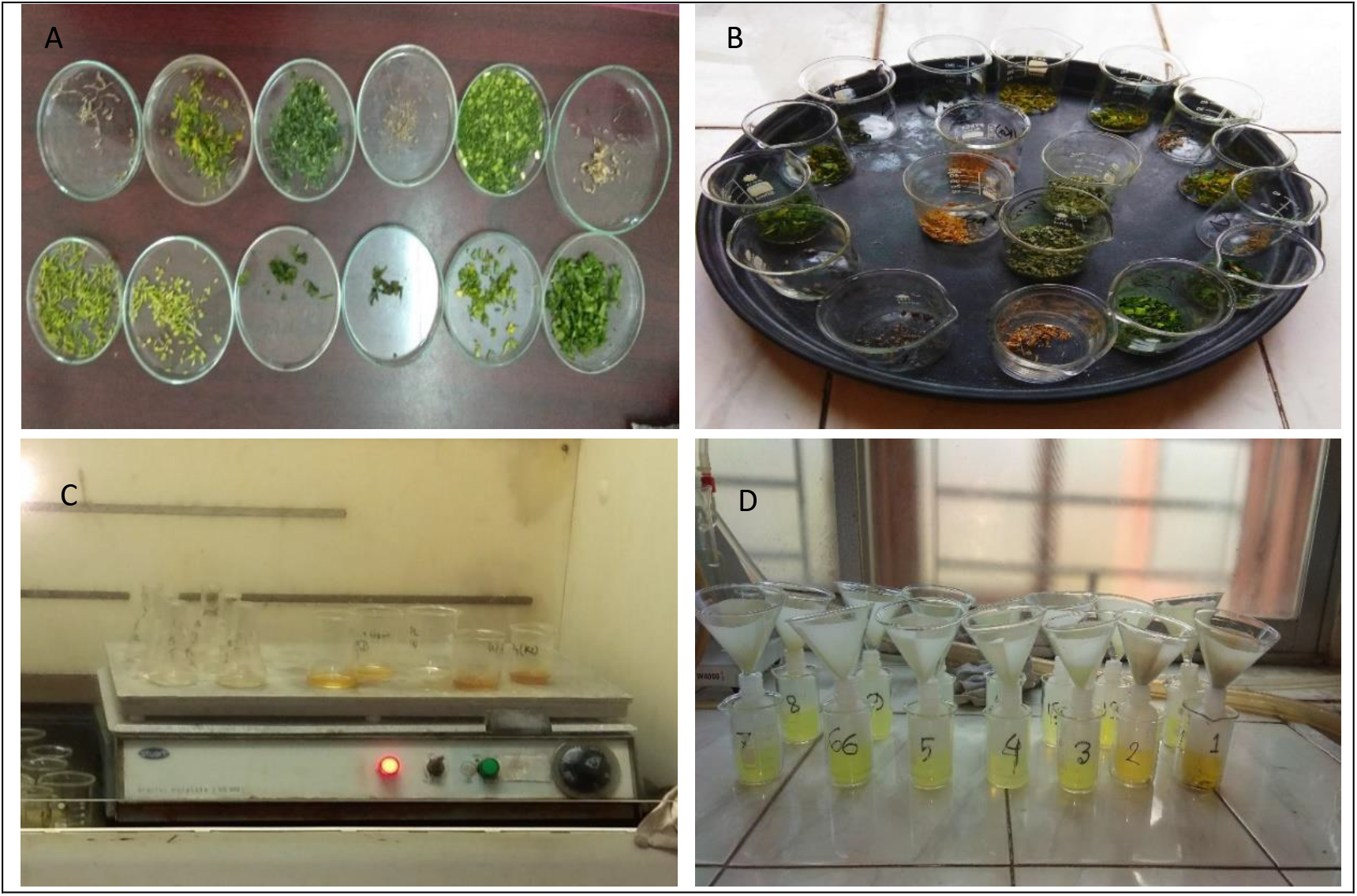
Preparation of samples for phytoremediation study: A) cutting of samples; B) weighted samples for metal analysis; C) slow boiling and evaporation process on hot plate; D) filtration of samples.

#### c) Measurement of heavy metal concentration by AAS

Three concentrations of standard solution of a particular metal were selected for metal analysis. After that, Blank solution was aspirated and adjusted to zero. A calibration curve was prepared for absorbance versus concentration of standard solution. The reading of the prepared sample solution was taken directly from the instrument. Heavy metal concentration was calculated by the following formulae:

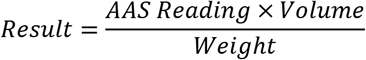

### Super-hyperaccumulator genotype selection and assessment of phytoremediation

After sowing the EMS induced seeds, plants were allowed to grow under heavy metal stress condition. Leaf samples were then assessed for phytoremediation. The EMS induced super-hyperaccumulator genotype was then selected. This EMS induced super-hyperaccumulator genotype produced seeds, which were collected for further planting. These next generation seeds were then sown in heavy metal polluted soil. Root, shoot, leaf and fruit of super-hyperaccumulator genotype and control *B. juncea* plant were then collected and phytoremediation study was performed as stated above. This experiment was conducted between February and May 2018.

### Statistical analysis

The concentration of Pb, Cr and Cd of 1-, 2-, and 3% EMS induced *B. juncea* were determined using three technical replicates of each sample, as only one best plant was used as a hyperaccumulator genotype after EMS and heavy metal treatments. In case of next generation super-hyperaccumulator genotype of *B. juncea* three biological replicates were used to assess phytoremediation. In all cases, mean value of technical/ biological replicates and their standard deviations were calculated using Microsoft Excel software.

## Results and discussion

It is very much expensive to remediate environmental pollutants by physico-chemical process. Therefore, using plants as a process to remediate soil pollutants is very much demanding now a days. Wild/cultivated plants those are used in phytoremediation are not able to accumulate heavy metals rapidly. In this regard, EMS induced hyperaccumulator plants may be the best alternative for this purpose. On the other hand, as *B. juncea* is considered as a suitable plant for phytoremediation, genetic approach to produce super phytoremediating genotype of *B. juncea* is also essential. EMS is a popular mutagenic organic compound that generates random mutations through nucleotide substitution/point mutations (Okagaski et al., 1991). In this experiment EMS was used in *B. juncea* seeds to develop hyperaccumulator genotype and then their phytoremediation potential was assessed in comparison to the control *B. juncea* plant.

### Selection of heavy metal tolerant plants

In this study, at first EMS treatment was carried out to generate heavy metal tolerant genotypes of *B. juncea*, which is considered as a hyperaccumulator plant in earlier studies (Choudhury et al., 2016; John et al., 2009). Heavy metal tolerant EMS induced plants of *B. juncea* was selected based on different parameters, such as seed germination rate, germination speed and plant survival rate.

#### Seed germination rate

For the calculation of seed germination rate of control and 1-, 2-, and 3% EMS induced *B. juncea* data was recorded and calculated at 7^th^ day of seed sowing. The highest seed germination rate (81.44%) was observed in the control *B. juncea* and the nearest (70.83) in 1% EMS induced *B. juncea* followed by 2-, and 3% EMS induced *B. juncea* (Table 1). It is also alarming that the lowest (36.84%) rate of seed germination is given by 3% EMS induced *B. juncea*, which revealed that this percentage of EMS is highly mutagenic than those of 1- and 2-% EMS, respectively.

**Table 1.**
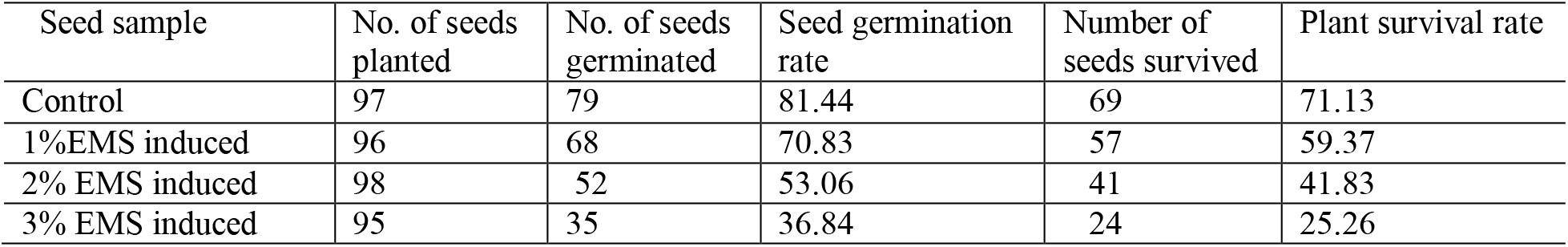
Comparative study of seed germination rate and plant survival rate

| Seed sample | No. of seeds planted | No. of seeds germinated | Seed germination rate | Number of seeds survived | Plant survival rate |
| --- | --- | --- | --- | --- | --- |
| Control | 97 | 79 | 81.44 | 69 | 71.13 |
| 1%EMS induced | 96 | 68 | 70.83 | 57 | 59.37 |
| 2% EMS induced | 98 | 52 | 53.06 | 41 | 41.83 |
| 3% EMS induced | 95 | 35 | 36.84 | 24 | 25.26 |

#### Plant survival rate

The highest (71.13%) plant survival rate was observed in control *B. juncea* and the nearest (59.37) was observed again in 1% EMS induced *B. juncea* (Table 1). On the other hand, 2- and 3% EMS induced *B. juncea* showed comparatively less plant survival rate. From the estimate of plant survival rate (Table 1) it is also evident that with the increase of mutagenicity plant survival rate was also decreased, which revealed that 2- and 3% EMS are not suitable to produce heavy metal hyper-accumulator genotypes.

#### Germination speed

For the calculation of germination speed of control and EMS induced genotypes, seeds were observed until 7^th^ day of seed sowing. Data were then recorded and calculated. As of seed germination rate and plant survival rate, similar results were found in case of seed germination speed (Table 2). The highest (24.05) germination speed was observed in control *B. juncea* and nearest (16.20) value was found in 1% EMS induced *B. juncea*. The lowest value of germination speed (8.6) was found in 3% EMS induced *B. juncea* that is approximately two times lower than 1% EMS induced *B. juncea*. By contrast, 2% EMS induced *B. juncea* has germination speed (12.65) in between 1- and 2-% EMS induced *B. juncea*.

**Table 2.** The seed germination speed of control, 1%, 2% and 3% EMS induced *Brassica juncea*

| Day of observation | Control |  |  | 1% |  |  | 2% |  |  | 3% |  |  |
| --- | --- | --- | --- | --- | --- | --- | --- | --- | --- | --- | --- | --- |
|  | n<sub>i</sub> | D<sub>i</sub> | GS | n<sub>i</sub> | D<sub>i</sub> | GS | n<sub>i</sub> | D<sub>i</sub> | GS | n<sub>i</sub> | D<sub>i</sub> | GS |
| 1<sup>st</sup> | 0 | 1 | 24.05 | 0 | 1 | 16.20 | 0 | 1 | 12.65 | 0 | 1 | 8.6 |
| 2<sup>nd</sup> | 0 | 2 |  | 0 | 2 |  | 0 | 2 |  | 0 | 2 |  |
| 3<sup>rd</sup> | 57 | 3 |  | 0 | 3 |  | 0 | 3 |  | 0 | 3 |  |
| 4<sup>th</sup> | 13 | 4 |  | 52 | 4 |  | 45 | 4 |  | 32 | 4 |  |
| 5<sup>th</sup> | 9 | 5 |  | 16 | 5 |  | 7 | 5 |  | 3 | 5 |  |
| 6<sup>th</sup> | 0 | 6 |  | 0 | 6 |  | 0 | 6 |  | 0 | 6 |  |
| 7<sup>th</sup> | 0 | 7 |  | 0 | 7 |  | 0 | 7 |  | 0 | 7 |  |

### Assessment of phytoremediation potential of EMS induced genotypes of *Brassica juncea*

From the comparative result (Table 1 & 2) it is obvious that with the increase of percentage of EMS seed germination rate, germination speed and plant survival rate were also decreased. Seed germination rate was used to determine EMS mutagenicity. Germination speed was used as an indication of the effectiveness of the plants to cope with heavy metal stress condition. Plant survival rate was used to determine heavy metal tolerance over 8- and 12-week planting period. As such, to select heavy metal hyperaccumulator super-genotype one from each of 1-, 2- and 3-% EMS induced eight-week-old plants named as BE21, BE11 and BE22 (Fig 3B), which showed better seed germination speed at 4^th^ day (the first day of seed germination); and a control plant, BC31 (Fig 3A) with seed germination speed at 3^rd^ day (the first day of seed germination for control group); with better plant survival capacity at 8-week planting period for both groups were selected for phytoremediation study. Phytoremediation was assessed with three technical replicates of each plant due to the selection of one heavy metal tolerant plant from each group.

**Fig 3.**
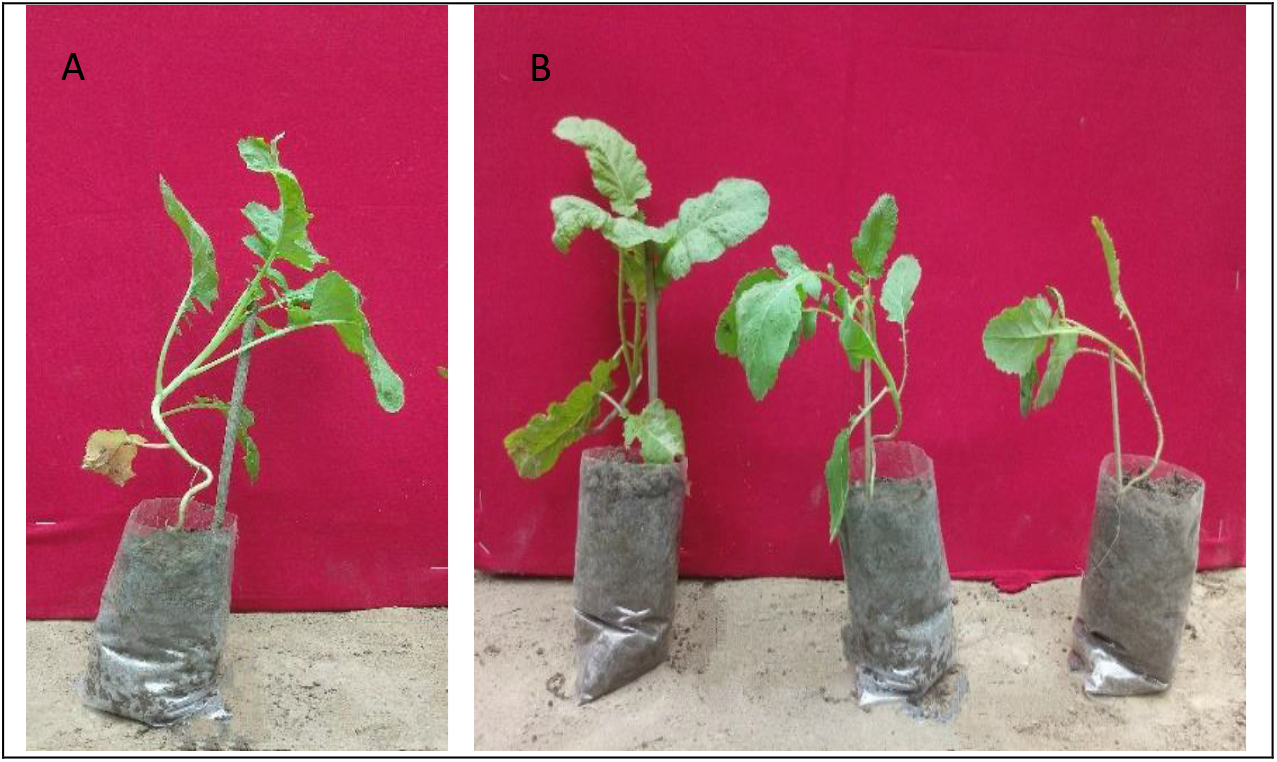
A) Control (BC31), and B) 1% (BE21), 2% (BE11) and 3% (BE22) EMS induced plants.

The highest concentration (33.36± 3.42 mg/kg dry weight) of Pb was accumulated by the leaf of 1% EMS induced BE21 genotype, which is four times more than the control plant (Table 3). In addition, 2- and 3% EMS induced genotypes accumulated less than 1% EMS induced genotype, but more than the control. In earlier study (Choudhury et al., 2016) Pb concentration in the leaf of *B. juncea* was 5.6 mg/kg dry weight and is six-fold less than the concentration found in BE21 genotype of current study, both at 8-week planting period.

**Table 3.** Comparative amount of Pb, Cr and Cd in the leaf of control and 1-, 2- and 3% EMS induced genotypes of *Brassica juncea* at 8-week planting period.

| Sample | Symbol | Weight (g) | Volume (ml) | Result (mg/kg dry weight) |  |  |
| --- | --- | --- | --- | --- | --- | --- |
|  |  |  |  | Pb | Cr | Cd |
| Surface soil | - | $0.50 \pm 0.01$ | 25 | $128.76 \pm 5.00$ | $117.03 \pm 4.51$ | $3.19 \pm 1.00$ |
| Control leaf | BC31 | $0.50 \pm 0.02$ | 25 | $8.01 \pm 1.00$ | $6.65 \pm 2.00$ | $0.24 \pm 0.08$ |
| 1% EMS induced leaf | BE21 | $0.50 \pm 0.01$ | 25 | $33.36 \pm 3.42$ | $7.34 \pm 2.94$ | $0.33 \pm 0.12$ |
| 2% EMS induced leaf | BE11 | $0.50 \pm 0.01$ | 25 | $28.60 \pm 2.01$ | $4.39 \pm 1.03$ | $0.33 \pm 0.13$ |
| 3% EMS induced leaf | BE22 | $0.50 \pm 0.01$ | 25 | $12.67 \pm 2.00$ | $5.42 \pm 1.98$ | $0.17 \pm 0.08$ |

It was found that 1% EMS induced leaf of BE21 genotype also accumulated the highest (7.34± 2.94 mg/kg dry weight) concentration of Cr; whereas 2-, and 3% EMS induced genotypes, BE11 and BE22, respectively accumulated less concentrations than 1% EMS induced genotype and the control. Previous research (Choudhury et al., 2016) found slightly more Cr concentration (8.4 mg/kg) in the leaf of *B. juncea*.

It was observed that accumulation of Cd in the leaf of BE21 (0.33± 0.12 mg/kg dry weight) and BE11 (0.33± 0.13 mg/kg dry weight) genotypes was similar. These concentration is highest in comparison to the control and approximately two-fold in comparison to the BE22 genotype. In a previous research (Zunaidi et al., 2021) Cd concentration in the leaf of *B. juncea* was 4.1±1.7 mg/kg at post-harvest period.

From this experiment it is concluded that 1% EMS induced genotype of *B. juncea* is the super-hyperaccumulator genotype. As a result, a further attempt was undertaken regarding the detail assessment of phytoremediation potential of *B. juncea* super-accumulator genotype obtained from this experiment.

### Assessment of phytoremediation potential of EMS induced next generation of *Brassica juncea* super-hyperaccumulator genotype

At maturity, seeds of the super-hyperaccumulator genotype, BE21 were collected and later on grown in the earthen pots filled with polluted Buriganga riverbank soil. Control *B. juncea* seeds were also grown separately in the earthen pots filled with polluted soil. After 12 weeks of seed germination, root, shoot, leaf and fruit (seeds) were collected and heavy metal phytoremediation was conducted. Three plants were selected randomly from control and super-hyperaccumulator genotype, respectively for assessment of phytoremediation.

### Comparative study of amount of Pb, Cr and Cd accumulated by next generation of *Brassica juncea* super-hyperaccumulator genotype

It was observed that the highest amount (91.53±6.59 mg/kg dry weight) of Pb was accumulated in the root of the next generation of *B. juncea* super-hyperaccumulator genotype, BE21. This concentration is approximately two-times higher than the control (Table 4) plant. Shoot and leaf of BE21 super-genotype accumulated approximately three-times higher concentration of Pb than the control, and fruit (seeds) accumulated slightly more concentration of Pb. The result of Pb concentration in the root, shoot and leaf of previous study (Choudhury et al., 2016) were 16.5, 6.0 and 6.4 mg/kg, respectively in 12-week period, which are much less than the super-hyperaccumulator genotype, BE21 of current research, at the same 12-week planting period. Previous another research (Payus and Talip, 2014) found Pb in the leaf and root of paddy rice as 0.26±0.14 and 7.70±1.27 mg/kg, respectively, which are much less than the present study. Other research (Liu et al., 2009) found Pb concentration in the root of wheat as 18.49–30.83 mg/kg, which is also less than the current study.

**Table 4.** Comparative amount of Pb, Cr and Cd in the root, shoot, leaf and fruit (seeds) of the control and next generation *Brassica juncea* super-hyperaccumulator genotype at 12-week planting period (n=3, ±=standard deviation).

| Sample | Weight (g) | Volume (ml) | Result (mg/kg dry weight) |  |  |
| --- | --- | --- | --- | --- | --- |
|  |  |  | Pb | Cr | Cd |
| Surface soil | $0.50 \pm 0.02$ | 25 | $162.47 \pm 9.02$ | $132.21 \pm 8.92$ | $3.30 \pm 1.92$ |
| Control root | $0.50 \pm 0.02$ | 25 | $53.60 \pm 4.91$ | $31.10 \pm 2.96$ | $0.90 \pm 0.17$ |
| Control shoot | $0.51 \pm 0.01$ | 25 | $10.68 \pm 1.01$ | $9.57 \pm 1.93$ | $0.39 \pm 0.02$ |
| Control leaf | $0.51 \pm 0.02$ | 25 | $9.07 \pm 1.00$ | $6.33 \pm 1.98$ | $0.39 \pm 0.12$ |
| Control fruit | $0.50 \pm 0.02$ | 25 | $4.79 \pm 0.81$ | $5.90 \pm 1.00$ | $0.10 \pm 0.01$ |
| Control total | - | - | 78.14 (48.10%) | 52.90 (40.01%) | 1.78 (53.94%) |
| BE21 root | $0.51 \pm 0.02$ | 25 | $91.53 \pm 6.59$ | $57.02 \pm 3.24$ | $1.96 \pm 0.92$ |
| BE21 shoot | $0.51 \pm 0.02$ | 25 | $33.31 \pm 1.01$ | $18.51 \pm 1.36$ | $0.19 \pm 0.01$ |
| BE21 leaf | $0.51 \pm 0.02$ | 25 | $28.35 \pm 3.61$ | $14.98 \pm 2.01$ | $0.52 \pm 0.32$ |
| BE21 fruit | $0.50 \pm 0.01$ | 25 | $5.59 \pm 0.93$ | $6.15 \pm 1.92$ | $0.19 \pm 0.02$ |
| BE21 total | - | - | 158.78 (97.73%) | 96.66 (73.11%) | 2.86 (86.67%) |

In the root of the next generation of BE21 genotype, the highest concentration of Cr was found (57.02±3.24 mg/kg dry weight), which is approximately two-fold higher than the control. Shoot of the next generation BE21 super-genotype accumulated two-times more Cr compared to the control. On the other hand, leaf of the next generation BES-21 super-genotype accumulated more than two-times Cr than the control. In the fruit (seeds), Cr content in the BE21 was also more than the control plant. The result of Cr concentration of root and shoot at 12-week planting period of previous study (Choudhury et al., 2016) were 61.55 and 30.0 mg/kg, respectively which are more, and in the leaf it was 11.0 mg/kg, which is less than BE21, the super-hyperaccumulator next generation genotype of current study. Another previous study (Payus and Talip, 2014) found much less Cr concentration in the root (5.46±2.26 mg/kg) and leaf (4.34±2.01 mg/kg) of rice plant than current research. In the root of wheat of earlier research (Liu et al., 2009) much less Cr concentration (2.55–8.07 mg/kg) was obtained than the present study.

Cd concentration was more in root (1.96±0.92 mg/kg dry weight), leaf (0.52±0.32 mg/kg dry weight) and fruit (0.19±0.02 mg/kg dry weight) and less in the shoot (0.19±0.01 mg/kg dry weight) of BE21 next generation super-hyperaccumulator genotype in comparison to the control plant. Earlier research (Gurajala et al., 2019) found Cd concentration of different *B. juncea* genotypes ranges between 0.44±0.03 and 2.94±0.05 mg/kg in the root, and between 0.31±0.02 and 2.43±0.00 mg/kg in the shoot at 90 days maturity period. Previous another research (Payus and Talip, 2014) conducted on rice found Cd concentration in the leaf and root as 0.11±0.05 mg/kg and 0.38±0.09 mg/kg, respectively— several folds less than the current research. By contrast, Cd concentration in the root (2.63–4.83mg/kg) of *T. aestivum* of other study (Liu et al., 2009) was more than current study.

From current study, it was investigated that concentration of Pb in the root of EMS induced *B. juncea* super-genotype, BE21 (91.53 mg/kg dry weight) was five times more than the result found by earlier study (Choudhury et al., 2016), which used *B. juncea* for the phytoremediation of Buriganga river sediment soil. Moreover, EMS derived BE21 super-genotype shoot uptakes more than five times Pb (33.31 mg/kg dry weight) and leaf uptakes approximately four times higher concentration (28.35 mg/kg dry weight) of Pb than the result found by earlier study (Choudhury et al., 2016). It revealed that EMS induced super-genotype has better heavy metal accumulation capacity compared to the non-EMS treated genotypes.

In earlier study (Choudhury et al., 2016), the average concentration of Pb and Cr in the Buriganga riverbed sediments were 34.9 mg/kg and 141.5 mg/kg, respectively and Pb and Cr uptakes were 28.9 mg/kg and 102.6 mg/kg, which are 83- and 73% accumulation of Pb and Cr, respectively when *B. juncea* was used at 12-week period (final growth phase).

In current study, it was observed that from root, shoot, leaf and fruit; root solely accumulated the highest concentration of these three types of heavy metals. In the root, concentration of Pb (91.53±6.59 mg/kg dry weight) of *B. juncea* super-genotype is approximately three times higher than the shoot, more than three times higher than the leaf, and approximately fifteen times higher than the fruit. Similarly, Cr and Cd concentration in the root were also several times higher than those found in the shoot, leaf and fruit of *B. juncea* super-genotype. The total concentration of Pb in the Buriganga riverbank surface soil was 162.47 mg/kg dry weight (Table 4). Root, shoot, leaf and fruit of *B. juncea* super-genotype accumulated a total of 158.78 mg/kg dry weight, which is equivalent to 98% Pb accumulation/phytoremediation. Similarly, total Cr and Cd accumulated by the root, shoot, leaf and fruit of *B. juncea* super-genotype were 73- and 87%, respectively. By contrast, total accumulation of Pb, Cr and Cd by the control (non-EMS treated) *B. juncea* plant is only 48-, 40- and 54%, respectively.

From these experiments it is revealed that EMS induced plants of *B. juncea* is suitable to produce hyperaccumulator genotypes. The most significant part of this research is that during the heavy metal stress condition in the field; as like root, shoot and leaf tissues; fruit (seeds) also uptakes heavy metal. Therefore, precaution should be taken accordingly while using plants for heavy metal phytoremediation in regard of using any edible plant part for human consumption or use them as fodder. Although different plants were used for phytoremediation of Buriganga riverbank soil of Bangladesh, however to the best of our knowledge, this is the first report of developing EMS induced hyperaccumulator genotypes of *B. juncea* for the same purpose.

## Competing interest

There is no competing interest regarding this study.

## Acknowledgement

The authors are grateful to Jagannath University and University Grant Commission of Bangladesh, respectively for providing some project assistance.

## Notes

### Competing Interest Statement

The authors have declared no competing interest.

